# Polycomb Cbx2 Condensates Assemble through Phase Separation

**DOI:** 10.1101/468926

**Authors:** Roubina Tatavosian, Samantha Kent, Kyle Brown, Tingting Yao, Huy Nguyen Duc, Thao Ngoc Huynh, Chao Yu Zhen, Brian Ma, Haobin Wang, Xiaojun Ren

## Abstract

Polycomb group (PcG) proteins are master regulators of development and differentiation. Mutation and dysregulation of PcG genes cause developmental defects and cancer. PcG proteins form condensates in the nucleus of cells and these condensates are the physical sites of PcG-targeted gene silencing. However, the physiochemical principles underlying the PcG condensate formation remain unknown. Here we show that Polycomb repressive complex 1 (PRC1) protein Cbx2, one member of the Cbx family proteins, contains a long stretch of intrinsically disordered region (IDR). Cbx2 undergoes phase separation to form condensates. Cbx2 condensates exhibit liquid-like properties and can concentrate DNA and nucleosomes. We demonstrate that the conserved residues within the IDR promote the condensate formation *in vitro* and *in vivo.* We further indicate that H3K27me3 has minimal effects on the Cbx2 condensate formation while depletion of core PRC1 subunits facilitates the condensate formation. Thus, our results reveal that PcG condensates assemble through liquid-liquid phase separation (LLPS) and suggest that PcG-bound chromatin is in part organized through phase-separated condensates.

## Introduction

The genome in eukaryotic cells can be broadly classified as euchromatin (active transcription) and heterochromatin (repression and silencing) (1,2). Heterochromatin can be further described broadly as constitutive and facultative heterochromatin (3-5). Constitutive heterochromatin is observed at and near centromeres and telomeres (5). Facultative heterochromatin is found at a specific subset of genes encoded for regulators of development and differentiation (3,4). Another example of facultative heterochromatin is X chromosome inactivation in female mammals (6). Facultative heterochromatin represses gene expression in part through compacting chromatin to reduce the accessibility of DNA (3-5). Facultative heterochromatin is decorated by the trimethylation of lysine 27 at histone H3 (H3K27me3), which is the catalytic product of Polycomb Repressive Complex (PRC) 2, one complex of Polycomb Group (PcG) proteins (3,4). H3K27me3 is the binding site for PRC1 (another complex of PcG proteins) that can compact chromatin, forming particular chromatin compartments (3,4). The repressed, compacted chromatin domains are preserved during cell division (3,4). These functions and behaviors of facultative heterochromatin have raised several fundamental questions. For example, what are the properties of facultative heterochromatin? How is compaction achieved? How do PcG proteins contribute to establishing facultative heterochromatin compartments? How do PcG proteins maintain facultative heterochromatin compartments?

Biochemical and genetic studies of PcG proteins have begun addressing some of the questions raised above. For example, in terms of understanding the properties of facultative heterochromatin, it has been shown that PcG proteins directly regulate chromatin structure and chemically modify the histones of facultative heterochromatin (7). PRC2 methylates histone H3 on lysine 27 (8-12). Eed, one core subunit of PRC2, binds H3K27me3, which allosterically stimulates PRC2 activity, and this stimulation can increase local H3K27me3 level (13,14). H3K27me3 provides a binding site for Cbx7 and Cbx8 of the Cbx family proteins (Cbx2/4/6/7/8) (15), which recruits canonical Cbx7-PRC1 and Cbx8-PRC1 to chromatin. PRC1 complexes can ubiquitinate histone H2A at lysine 119 (H2AK119Ub) (16,17), which can stimulate PRC2 activity (18,19). Thus, a feedback loop between PRC1 and PRC2 is created to reinforce the epigenetic modifications of facultative heterochromatin. In terms of understanding compaction, it has been demonstrated that the compaction function of PRC1 in mammals is facilitated by Cbx2 of the Cbx family proteins (20,21). Mutation of the Cbx2 residues that are required for compaction leads to homeotic transformations that are similar to those observed with PcG loss-of-function mutations (21). In terms of understanding facultative heterochromatin compartments, it has been documented that, in the nucleus, PcG proteins form microscopically visible condensates (22-26). PcG condensates function as specific nuclear compartments for target gene silencing (22-26). Overall, these biochemical and genetic studies suggest that PRC1 and PRC2 coordinate to establish and maintain facultative heterochromatin. Despite these exciting advances, the fundamental physicochemical principles that underpin how PcG proteins establish, maintain, and regulate facultative heterochromatin remain incompletely understood.

Spatial and temporal compartmentalization of intracellular components into organelles in eukaryotic cells is a generic theme for organizing biochemical reactions (27-32). These organelles can be membrane-bound or membraneless. A large number of membraneless compartments, including the nucleolus, stress granules, Cajal bodies, PML nuclear bodies, and others, are condensates formed by condensation of cellular components through liquid-liquid phase separation (LLPS) (27-31). The forces that drive LLPS are multivalent interactions among proteins and other macromolecular polymers such as RNA and DNA (27-29). Phase-separated condensates have been shown to be involved in multiple cellular processes and functions (27-29). Over the past year, phase-separated membraneless condensates have been suggested to be implicated in transcriptional activation and repression (33-39). A phase-separated model has been emerging to explain transcriptional activation: transcription factors and coactivators phase separate to form condensates that interact with condensates of RNA polymerase II (RNA Pol II) to efficiently activate transcription (33-37). Phase-separated condensates also function in transcriptional repression. Heterochromatin protein 1α (HP1α) phase separates to form condensates that compartmentalize constitutive heterochromatin (38,39). Facultative heterochromatin represents one major class of chromatin structures. Whether the PcG proteins that are responsible for the formation of facultative heterochromatin phase separate to form condensates remains unknown.

Here we provide the first experimental evidence that the PRC1 protein Cbx2 phase separates to form condensates that can concentrate DNA and nucleosome. The conserved residues within the Cbx2 IDR are required for the Cbx2 condensate formation *in vitro* and *in vivo.* We show that H3K27me3 contributes little to the Cbx2 condensate formation, while depletion of Cbx2-PRC1 subunits facilitates the condensate formation. Thus, our results provide a general experimental framework that can explain how PcG condensates assemble and a starting point for further exploring how phase separation facilitates efficient and specific control of transcription.

## Results

### Cbx2 forms condensates in cells

We first investigated whether Cbx2 forms condensates in living cells. We integrated *YFP*-*Cbx2* and *HaloTag (HT)*-*Cbx2* into the genome of PGK12.1 mouse embryonic stem (wild-type mES) cells, respectively. To observe the cellular distribution of HT-Cbx2, we labelled the fusion protein by HaloTag TMR ligand in living cells. Both YFP-Cbx2 and HT-Cbx2 formed condensates in living wild-type mES cells (**Fig. 1a-b**), which is consistent with the previous observations reporting that both exogenous and endogenous Cbx2 forms condensates (40,41). About half of cells contained microscopically visible Cbx2 condensates. The average area of condensates was 0.19 μm^2^ (~250 nm in radius), and their fluorescence intensity was ~1.5-fold higher than the average intensity. It is possible that the formation of Cbx2 fusion condensates in mES cells is due to their overexpression. To resolve this possibility, we integrated *YFP*-*Cbx2* and *HT*-*Cbx2* into the genome of *Cbx2*^−/−^ mES cells. The distribution of YFP-Cbx2 and HT-Cbx2 in *Cbx2*^−/−^ mES cells was similar to that in wild-type mES cells (**Fig. 1a-b**). These data indicate that Cbx2 forms microscopically visible condensates in living cells.

**Figure 1.**
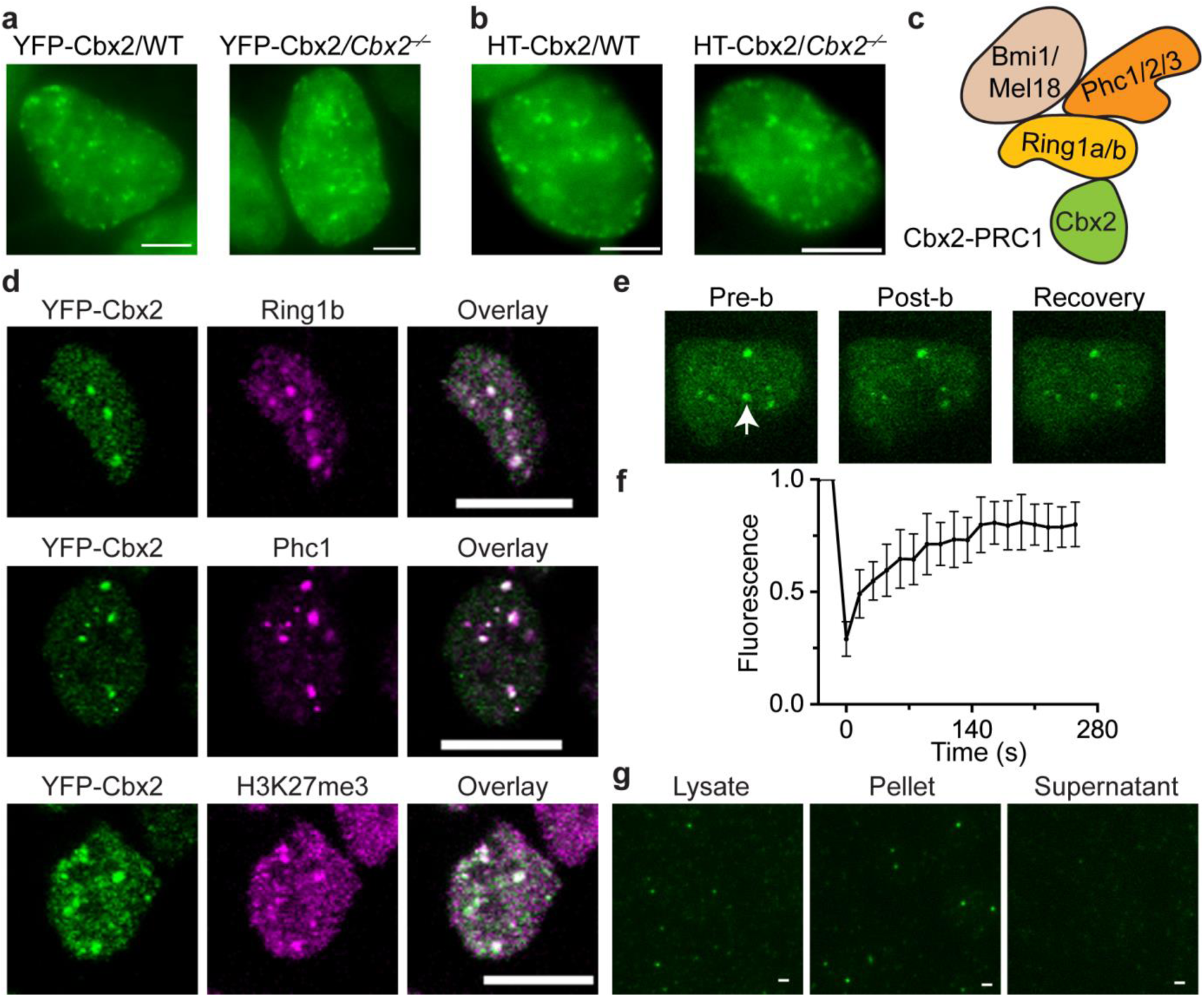
Cbx2 phase separates to form condensates in cells. a. Live-cell epi-fluorescence images of YFP-Cbx2 in wild-type (WT) and *Cbx2*^−/−^ mES cells. Scale bar, 5.0 μm.
b. Live-cell epi-fluorescence images of HaloTag (HT)-Cbx2 in wild-type and *Cbx2*^−/−^ mES cells. HT-Cbx2 was labelled with HaloTag TMR ligand. Scale bar, 5.0 μm.
c. Schematic representation of the core subunits of the Cbx2-PRC1 complex.
d. Confocal images of immunostained YFP-Cbx2 and H3K27me3 in wild-type mES cells as well as endogenous Ring1b and Phc1. The cells were stained by using antibodies against YFP (green), Ring1b (magenta), Phc1 (magenta), and H3K27me3 (magenta). Overlay images are shown. Scale bar, 10 μm.
e. Representative FRAP images of YFP-Cbx2 stably expressed in wild-type mES cells. The images were taken before (Pre-b) and after (Post-b) photobleaching. The condensate that was bleached is indicated by white arrowhead.
f. FRAP curve of YFP-Cbx2 in wild-type mES cells that stably express YFP-Cbx2. The FRAP curve is obtained from averaging data from ten cells.
g. Epi-fluorescence imaging of YFP-Cbx2 condensates isolated from cells. Cells stably expressing YFP-Cbx2 were cross-linked with formaldehyde. Lysate was prepared. Both lysate and resuspended pellets contained YFP-Cbx2 condensates; however, supernatant did not have condensates. Scale bar, 2.0 μm.

Cbx2 forms a stable PRC1 complex (Cbx2-PRC1), including Phc1 and Ring1b (**Fig. 1c**) (42), so we investigated whether YFP-Cbx2 condensates colocalize with Cbx2-PRC1 subunits. We stained endogenous Ring1b and Phc1 as well as YFP-Cbx2. Ring1b and Phc1 formed condensates in cells (**Fig. 1d**), consistent with the previous reports (23,41,43,44). YFP-Cbx2 condensates colocalized with condensates of Ring1b and Phc1 (**Fig. 1d**). Since H3K27me3 marks PcG-targeted genes, we investigated whether Cbx2 condensates colocalize with H3K27me3. Immunofluorescence of H3K27me3 and YFP-Cbx2 showed that Cbx2 condensates colocalize with chromatin with the dense H3K27me3 mark (**Fig. 1d**), suggesting that PcG-targeted genes are recruited to Cbx2 condensates, or vice versa. Thus, our results show that Cbx2 condensates colocalize Cbx2-PRC1 subunits and H3K27m3-marked chromatin regions.

Next, we interrogated whether Cbx2 condensates exhibit liquid-like features that are characterized with rapid exchange kinetics, which can be studied by measuring the recovery rate using fluorescence recovery after photobleaching (FRAP). We performed FRAP experiments on condensates of YFP-Cbx2 stably expressed in mES cells. FRAP showed that 80% of YFP-Cbx2 within condensates is recovered within 3 min (**Fig. 1e-f**), consistent with our previous reports (40). These results indicate that Cbx2 within condensates dynamically exchange with surrounding environments and has liquid-like properties in cells.

If Cbx2 condensates are liquid-like, reducing concentration of YFP-Cbx2 would dissolve these condensates. We lysed cells stably expressing YFP-Cbx2 in lysis buffer to cause a local decrease of YFP-Cbx2 through diffusion. We did not detect YFP-Cbx2 condensates in the lysate. We expected that formaldehyde crosslinking would preserve Cbx2 condensates. After sonication and centrifugation, YFP-Cbx2 condensates would be within the pellets. To test this speculation, prior to lysis, we cross-linked cells with formaldehyde. Lysates were prepared from the cross-linked cells and subjected to sonication. Using fluorescence microscopy, we observed Cbx2 condensates within the sonicated lysate (Fig. 1g). After centrifugation, we did not observe Cbx2 condensates in the supernatant, but instead in the re-suspended pellets (Fig. 1g). These data further support that Cbx2 condensates form in the cells and possess liquid-like properties.

### Cbx2 phase separates to form condensates *in vitro*

Proteins can undergo LLPS through intrinsically disordered regions (IDRs), leading to condensate formation in aqueous solution (27-29). We analyzed the properties of the primary sequence of Cbx2 and found that 59% of the Cbx2 sequence is intrinsically disordered as predicted by MobiDB 3 (**Fig. 2a**) (45). Thus, we reasoned that Cbx2 could form condensates *in vitro* through LLPS. To test this hypothesis, we expressed and purified recombinant GST-Cbx2-FLAG (GST-Cbx2) from *E. coli* at high salt concentration or in the presence of glutathione (GSH) (**Fig. 2b**). We found that both high salt and glutathione prevent aggregating of Cbx2. We placed the tags at the respective N-terminal and C-terminal ends of Cbx2 to remove truncated Cbx2 during the purification. We dialyzed the high salt of GST-Cbx2 fusion to 140 mM NaCl at 4 °C overnight and transferred 10 μl of sample to coverslip. After condensates settled down on the surface of coverslip, we performed differential-interference-contrast (DIC) imaging and observed Cbx2 condensates with a size of a few hundred nanometers (**Fig. 2c**). Next, we generated Cbx2-FLAG (Cbx2) without GST fusion (**Fig. 2b**). Cbx2 also underwent LLPS to form condensates (**Fig. 2c**), suggesting that the condensate formation is not driven by GST. To determine the identity of these condensates, we produced GST-YFP-Cbx2-FLAG (GST-YFP-Cbx2) (**Fig. 2b**). Fluorescence imaging showed that GST-YFP-Cbx2 assemble to condensates (**Fig. 2c**). Under the same conditions, GST and BSA did not form condensates (**Fig. 2c**). LLPS typically depends on the concentration of components in the system, so we performed the condensate formation assay with varying concentrations of Cbx2, ranging from 1.2 μM to 12 μM. We found that the phase separation of Cbx2 is concentration-dependent (**Fig. 2d**). Thus, our results demonstrate that Cbx2 can undergo LLPS to form condensates *in vitro.*

**Figure 2.**
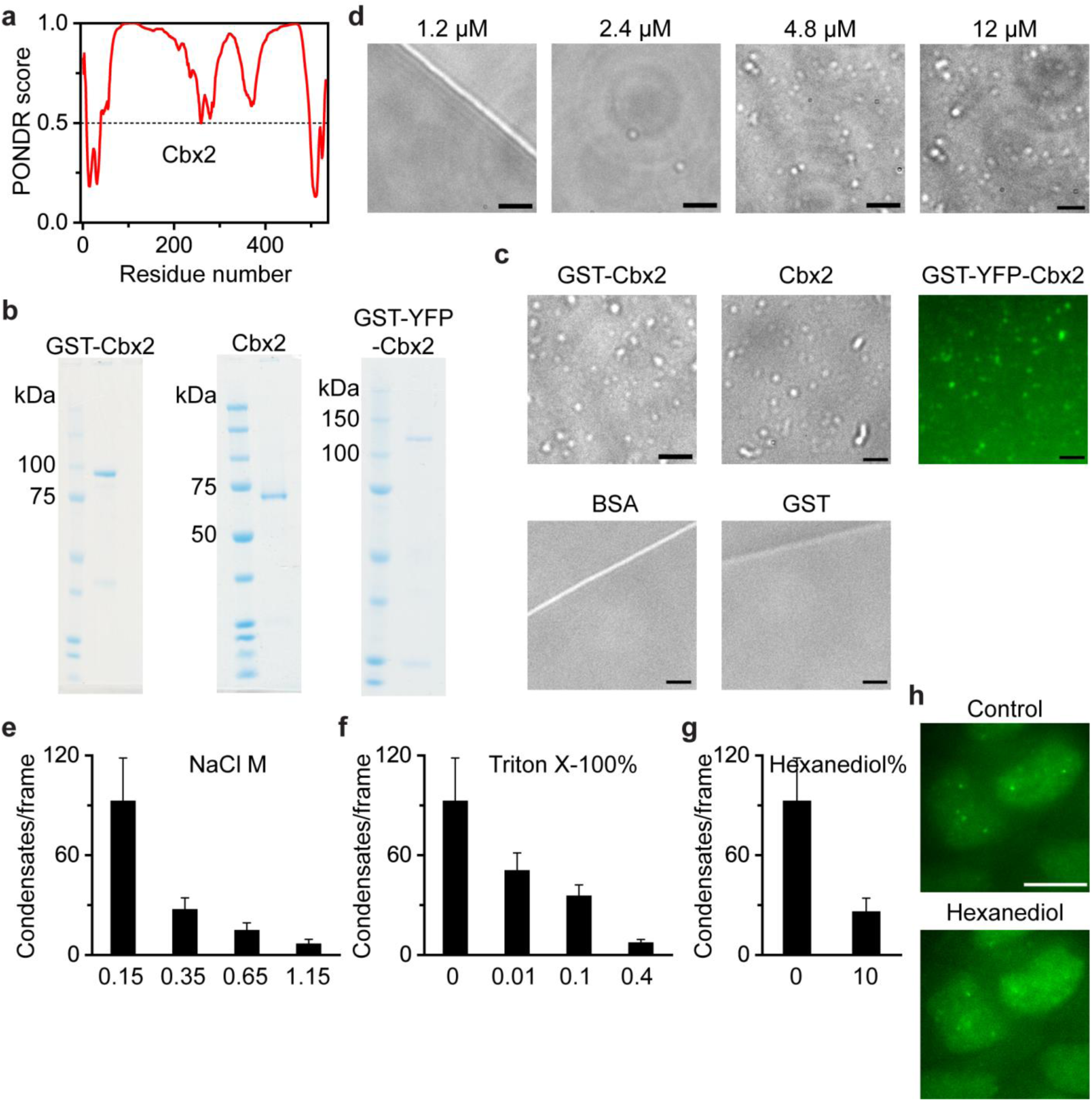
Cbx2 phase separates to form condensates *in vitro*. a. Cbx2 is an intrinsically disordered protein predicted by MobiDB 3 (45). The PONDR score being greater than 0.5 indicates intrinsically disordered regions. 59% of Cbx2 sequence is intrinsically disordered.
b. SDS-PAGE analysis of recombinant Cbx2 proteins. Left: recombinant GST-Cbx2-FLAG (GST-Cbx2). Middle: recombinant Cbx2-FLAG (Cbx2). Right: recombinant GST-YFP-Cbx2-FLAG (GST-YFP-Cbx2). Molecular weight ladder is shown at the left of the gel image.
c. Representative differential-interference-contrast (DIC) images of GST-Cbx2 and Cbx2 condensates as well as the control BSA and GST on the surface of coverslip. GST-YFP-Cbx2 condensates are representative epi-fluorescence image. Scale bar, 2.0 μm.
d. Dependence of the formation of Cbx2 condensates on its concentrations. Representative DIC images of condensates on the surface of coverslip are shown. Scale bar, 2.0 μm.
e. Increasing concentration of NaCl, Triton X-100, and hexanediol dissolves Cbx2 condensates. Cbx2 condensates were incubated with indicated concentrations of NaCl, Triton X-100, and hexanediol for 30 min on ice. The mixture was loaded to coverslip for imaging. Condensates then were counted by using ImageJ.
f. Representative epi-fluorescence images of wild-type mES cells stably expressing YFP-Cbx2 before and after treatment with 10% hexanediol for 5.0 min. Scale bar, 10 μm.

Classical polymer theory predicts that polymers undergo LLPS through multivalency-driven interactions such as cation-pi, electrostatic, dipolar, and hydrophobic interactions (27-29). Thus, we investigated whether NaCl and Triton X-100 dissolve Cbx2 condensates. We found that treatment of Cbx2 condensates with increasing concentrations of NaCl and Triton X-100 causes a reduction in the number of Cbx2 condensates (**Fig. 2e-f**). Hexanediol is known to dissolve liquid-like condensates (33,35), possibly through disruption of hydrophobic interactions. We found that treatment of Cbx2 condensates *in vitro* as well as in mES cells expressing YFP-Cbx2 with Hexanediol results in a reduction in the number of condensates (**Fig. 2g-h**). These results indicate that multivalent interactions contribute to the LLPS of Cbx2.

### Cbx2 condensates concentrate DNA and nucleosomes

One of the characteristic properties of cellular compartmentalization is that compartments can increase the local concentration of resident biochemical molecules (27-32). PcG condensates are the physical sites of PcG-mediated silencing, involving organization of the PcG-bound chromatin (22-24). Given that Cbx2 can directly bind DNA (46), we investigate whether Cbx2 condensates can concentrate DNA. We labelled 24-bp double-stranded DNA with fluorescent dye and mixed them with Cbx2. Dye-labelled DNA did not form condensates; however in the presence of Cbx2, DNA was concentrated and the concentrated DNA was colocalized with Cbx2 condensates (**Fig. 3a**). Previous studies have shown that Cbx2 can compact chromatin on its own (20), so we tested whether Cbx2 can concentrate core nucleosome particles. We prepared dye-labelled nucleosomes and mixed them with Cbx2. After dialysis, we observed that Cbx2 condensates colocalize with the concentrated dye-labelled nucleosomes (**Fig. 3a**). Under the same conditions, Cbx2 condensates could not enrich dye (**Fig. 3a**). Thus, these results suggest that Cbx2 condensates can concentrate DNA and nucleosomes *in vitro.*

**Figure 3.**
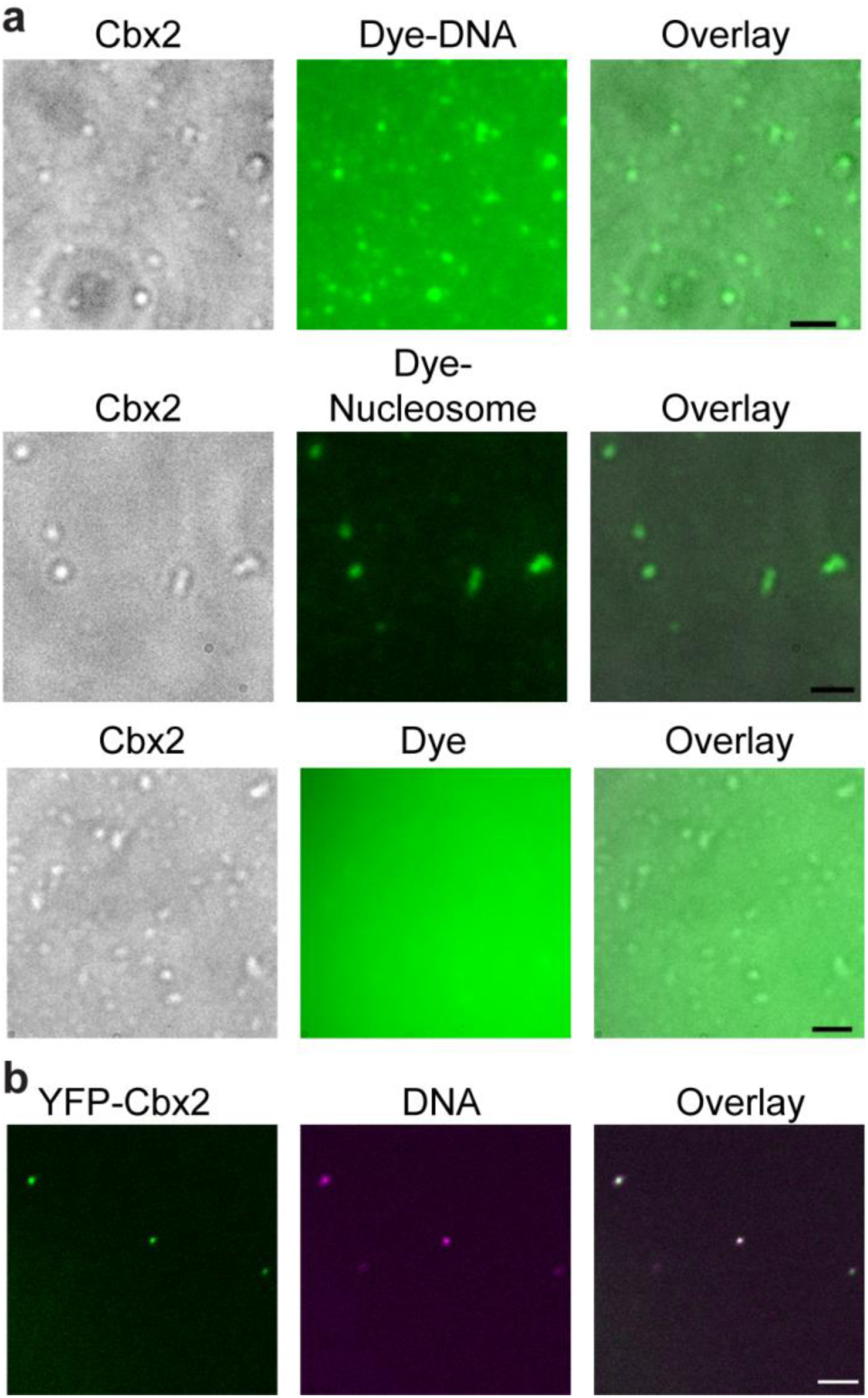
Cbx2 condensates concentrate DNA and nucleosomes. a. Micrographs of phase-separated Cbx2 condensates with Alexa 488, Alexa 488-labelled DNA, or Cy5-labelled nucleosome. Cbx2 condensates are DIC images on the surface of coverslip. Alexa 488, Alexa 488-labelled DNA, and Cy5-labelled nucleosome are fluorescence images. Overlay images are shown. Scale bar, 2 μm.
b. Epi-fluorescence imaging of DNA and YFP-Cbx2 condensates isolated from cells. Cells stably expressing YFP-Cbx2 were cross-linked with formaldehyde. Lysate was prepared. Resuspended pellets were stained with Hoechst. Overlay image is shown. Scale bar, 5.0 μm.

Given that PcG condensates are the repressive compartments for PcG-targeted genes, we should be able to detect DNA with Cbx2 condensates isolated from cells. We cross-linked cells stably expressing YFP-Cbx2 with formaldehyde. After sonication and centrifugation, we resuspended the pellet and stained DNA with Hoechst. Fluorescence images showed that YFP-Cbx2 condensates contain concentrated DNA labelled by Hoechst (**Fig. 3b**). Our data indicate that Cbx2 condensates can enrich chromatin/DNA within cells.

### Conserved residues within the IDR of Cbx2 contribute to LLPS *in vitro* and in living cells

The driving forces of phase separation of proteins containing IDRs are non-covalent interactions, particularly cation-pi and electrostatic interactions (27-32). The cation-pi interactions occur between aromatic residues with Lys or Arg residues (47-52). We found that Cbx2 contains 48 Lys (9.0%) and 33 Arg (6.2%), whose frequency is higher than their respective average frequency in vertebrate proteins (6.6% for Lys and 4.9% for Arg) (53). There are 5 Phe (0.9%), 5 Trp (0.9%), and 3 Tyr (0.6%) in Cbx2, whose frequency is lower than their respective average frequency in vertebrate proteins (3.6% for Phe, 1.3% for Trp, and 3.4% for Tyr) (53). Given that proteins whose phase separations are promoted by the cation-pi interactions contain a high content of aromatic residues (47), we focused on the distribution of charged residues of Cbx2. Electrostatic interaction is one of the major driving forces that promote the phase separation of IDR-containing proteins (39,51,54-58). In Cbx2, many positively and negatively charged residues are grouped into a series of clusters across the sequence (**Fig. 4b**). It is interesting to note that the three conserved regions, AT-hook (ATH), ATH-like 1 (ATHL1), and ATHL2 (59), are positively charged clusters (**Fig. 4a-b**). We substituted residues PRG (77-79) for AAA to generate Cbx2^ATH^, PRG (134–136) for AAA to generate Cbx2^ATHL1^, and RKKRGRK (161–167) for AAAAGAA to generate Cbx2^ATHL2^ (**Fig. 4b**). The net positive charge per residue of the mutated regions of Cbx2^ATH^ and Cbx2^ATHL1^ was slightly reduced compared to Cbx2, while the net positive charge per residue of the mutated region of Cbx2^ATHL2^ was completely eliminated (**Fig. 4b**). We generated these mutant proteins and compared their ability to form condensates with Cbx2 *in vitro.* Our analysis indicated that the phase-separation ability of the three mutants is greatly reduced compared to Cbx2 (**Fig. 4c-d**). Cbx2^ATH^ and Cbx2^ATHL1^ had a better capacity to phase separate than Cbx2^ATHL2^ (**Fig. 4c-d**), consistent with the complete loss of positive charge in the ATHL2 of Cbx2^ATHL2^. Within the IDR of Cbx2, there is a conserved serine-rich region (SRR) consisting of a stretch of consecutive 19 residues of serine and threonine (59). We substituted SKSKSSSSSSSSTSSSSSS (102–120) for SKSKASASASASTASASAA to generate Cbx2^SRR^ (**Fig. 4a**). Cbx2^SRR^ greatly reduced its ability to phase separate compared to Cbx2 (**Fig. 4c-d**). Thus, these data indicate that these conserved residues within the IDR of Cbx2 promote LLPS *in vitro.*

**Figure 4.**
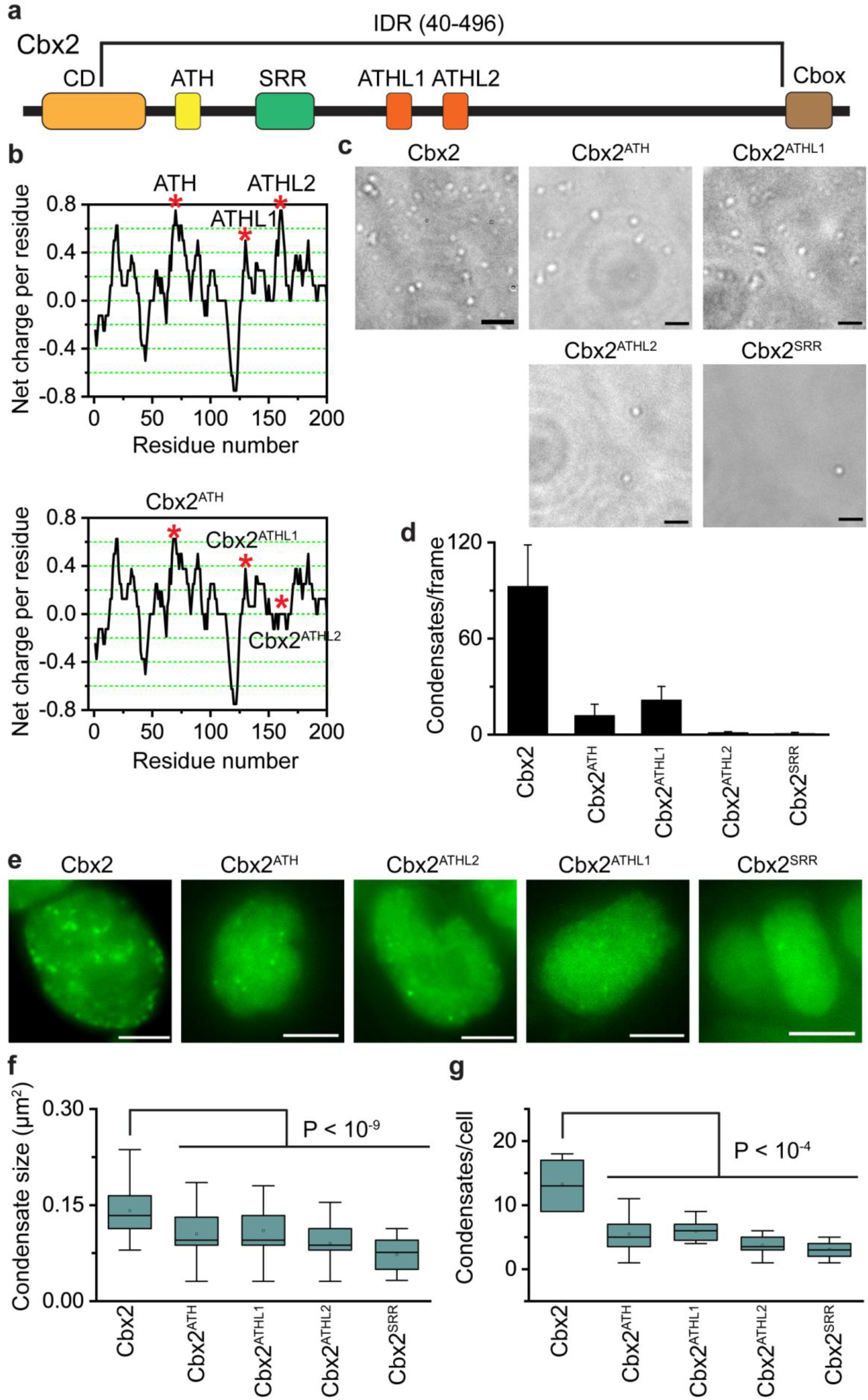
Conserved residues promote the LLPS of Cbx2. a. Schematic representation of Cbx2. IDR, intrinsically disordered region predicted by MobiDB 3 (45) (**Fig. 2a**). Conserved regions are shown: CD, chromodomain; ATH, AT-hook; ATHL1, AT-hook-like 1; ATHL2, AT-hook-like 2; and Cbox, Chromobox (59).
b. Charge distribution of Cbx2 and its mutants calculated by EMBOSS charge. The net charge per residue was averaged over a sliding window of eight residues. The three covered regions, ATH, ATHL1, and ATHL2, are positively charged (top panel). The three conserved regions were mutated, respectively, to generate Cbx2^ATH^, Cbx2^ATHL1^, and Cbx2^ATHL2^ (bottom panel).
c. Representative DIC images of condensates of Cbx2 and its variants (Cbx2^ATH^, Cbx2^ATHL1^, Cbx2^ATHL2^, and Cbx2^SRR^) on the surface of coverslip. The formation of condensates was performed under the same conditions.
d. Quantification of condensates per frame from **Fig. 4c**. Results are means ± SD.
e. Representative epi-fluorescence images of wild-type mES cells that stably express HT-Cbx2 and its variants fused with HT (Cbx2^ATH^, Cbx2^ATHL1^, Cbx2^ATHL2^, and Cbx2^SRR^), respectively. HT-Cbx2 and HT-Cbx2 variants were labelled by HaloTag TMR ligand, and live-cell epifluorescence imaging was performed. Scale bar, 5.0 μm.
f. Box plot of the condensate sizes for HT-Cbx2 and HT-Cbx2 variants (Cbx2^ATH^, Cbx2^ATHL1^, Cbx2^ATHL2^, and Cbx2^SRR^) in living mES cells. Data were obtained from at least 10 cells. P-value was calculated based on student’s t-test.
g. Box plot of the number of condensates for HT-Cbx2 and HT-Cbx2 variants (Cbx2^ATH^, Cbx2^ATHL1^, Cbx2^ATHL2^, and Cbx2^SRR^). Data were obtained from at least 10 cells. P-value was calculated based on student’s t-test.

To investigate whether these conserved residues of Cbx2 contribute LLPS *in vivo,* we established wild-type mES cells stably expressing *HT*-*Cbx2*^*ATH*^, *HT*-*Cbx2^ATL1^*, *HT*-*Cbx2^ATL2^*, or *HT*-*Cbx2^SRR^*. These Cbx2 mutants were labelled with HaloTag TMR ligand. We performed live-cell imaging of these mutants (**Fig. 4e**). Quantitative analysis showed that the size and the number of condensates of these Cbx2 mutants are significantly reduced compared to wild-type Cbx2 (**Fig. 4f-g**). We also noted that the size and the number of condensates of Cbx2^ATHL2^ and Cbx2^SRR^ were slightly smaller than Cbx2^ATH^ and Cbx2^ATHL1^ (**Fig. 4f-g**), consistent with *in vitro* analysis. Thus, our data demonstrate the conserved residues within the IDR that are critical for the LLPS of Cbx2 *in vitro* are also critical for the formation of Cbx2 condensates *in vivo.*

### H3K27me3 has minimal effects on the formation of Cbx2 condensates

The PRC2-catalyzed product H3K27me3 is the marker of PcG-targeted chromatin (7). H3K27me3 has been hypothesized to be the mark for recruiting Cbx-PRC1 to chromatin (**Fig. 5a**) (60). Thus, we asked whether H3K27me3 affects the formation of Cbx2 condensates *in vivo.* To this end, we integrated *HT*-*Cbx2* into the genome of *Eed*^−/−^ mES cells. Eed is the core component of PRC2 and *Eed* knockout results in a complete loss of H3K27me3 (15). Live-cell imaging of HT-Cbx2 labelled with HaloTag TMR ligand showed that HT-Cbx2 forms condensates in *Eed*^−/−^ mES cells (**Fig. 5c**). Quantitative analysis indicated that the size and the number of Cbx2 condensates in *Eed*^−/−^ mES cells are not significantly different from that in wild-type mES cells (**Fig. 5d-e**). Previous studies have shown that the aromatic cage, consisting of three aromatic residues, of the chromodomain (CD) domain of Cbx2 is critical for the H3K27me3 binding *in vitro* (61). We mutated the cage residue Phe-12 of Cbx2 to Ala (Cbx2^F12A^) (**Fig. 5b**). We stably expressed *HT*-*Cbx2^F12A^* in wild-type mES cells. Live-cell imaging showed that HT-Cbx2^F12A^ forms condensates (**Fig. 5c**). The size and the number of HT-Cbx2^F12A^ were similar to HT-Cbx2 (**Fig. 5d-e**). These data suggest that H3K27me3 contributes little to the formation of Cbx2 condensates in living cells.

**Figure 5.**
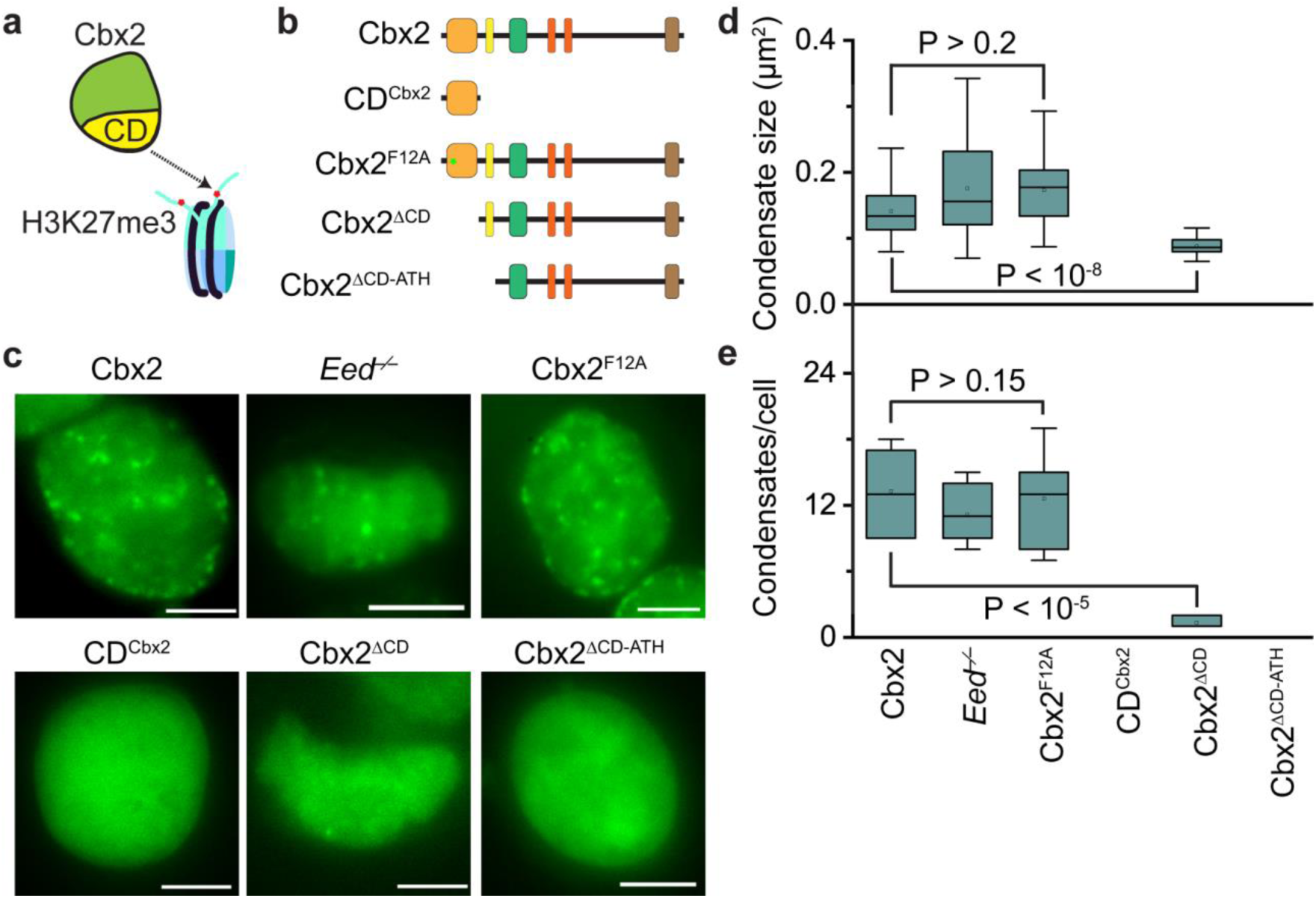
H3K27me3 has minor effects on the Cbx2 condensate formation. a. A hypothetic model for targeting Cbx2 to chromatin. It has been proposed that Cbx2 is recruited to chromatin through the interactions between CD and H3K27me3.
b. Schematic representation of Cbx2 and its variants used in the study. CD is the binding domain for H3K27me3 *in vitro.* The residue Phe-12 is the key one involved in the H3K27me3 binding *in vitro.*
c. Representative epi-fluorescence images for HT-Cbx2 in wild-type mES cells replicated from **Fig. 1b**, for HT-Cbx2 in *Eed*^−/−^ mES cells, and for HT-Cbx2^F12A^, HT-CD^Cbx2^, HT-Cbx2^ΔCD^, and HT-Cbx2^ΔCD-ATH^ in wild-type mES cells. HT-Cbx2 fusions were labelled by HaloTag TMR ligand, and live-cell epi-fluorescence imaging was performed. Scale bar, 5.0 μm.
d. Box plot of the condensate sizes for HT-Cbx2 in wild-type mES cells replicated from **Fig. 4f**, for HT-Cbx2 in *Eed*^−/−^ mES cells, and for HT-Cbx2^F12A^, HT-CD^Cbx2^, HT-Cbx2^ΔCD^, and HT-Cbx2^ΔCD-ATH^ in wild-type mES cells. Data were obtained from at least 10 cells. P-value was calculated based on student’s t-test.
e. Box plot of the number of condensates for HT-Cbx2 in wild-type mES cells replicated from **Fig. 4g**, for HT-Cbx2 in *Eed*^−/−^ mES cells, and for HT-Cbx2^F12A^, HT-CD^Cbx2^, HT-Cbx2^ΔCD^, and HT-Cbx2^ΔCD-ATH^ in wild-type mES cells. Data were obtained from at least 10 cells. P-value was calculated based on student’s t-test.

Since the CD of Cbx2 is the binding domain for H3K27me3 *in vitro* (61,62) (**Fig. 5a**), we investigated the effects of the CD on the formation of Cbx2 condensates in living cells. We fused CD with HT, generating HT-CD^Cbx2^ (**Fig. 5b**). We also deleted CD to generate HT-Cbx2^ΔCD^ (**Fig. 5b**). We established mES cells that stably express *HT*-*CD^Cbx2^* and *HT*-*Cbx2^ACD^*, respectively. HT-CD^Cbx2^ did not form condensates in living cells (**Fig. 5c**). HT-Cbx2^ΔCD^ phase separated to form condensates (**Fig. 5c**); however, their size and number significantly reduced in comparison with HT-Cbx2 (**Fig. 5d-e**). Given that ATH is adjacent to CD and binds DNA (46) (**Fig. 5b**), we deleted both CD and ATH to generate *HT*-*Cbx2*^Δ*CD*-*ATH*^ (**Fig. 5b**), which was then stably integrated into the genome of mES cells. HT-Cbx2^ΔCD-ATH^ did not phase separate to form condensates within living cells (**Fig. 5c**). Thus, these results indicate that the interactions of H3K27me3 and Cbx2 are not required for the formation of Cbx2 condensates; however, the amino acid residues within CD are required for the condensate formation.

### Depletion of Cbx2-PRC1 subunits facilitates the formation of Cbx2 condensates

Cbx2 phase separates on its own *in vitro,* so we speculate that removal of Cbx2-PRC1 subunits would not prevent the formation of Cbx2 condensates *in vivo.* To address this, we integrated *HT*-*Cbx2* into the genome of *Ring1a*^−/−^*/Ring1b^fl/fl^*; *Rosa26*::CreERT2 and *Bmi1*^−/−^*/Mei18*^−/−^ mES cells, respectively. *Ring1b* in *Ring1a*^−/−^*/Ring1b^fl/fl^*; *Rosa26*::CreERT2 mES cells was depleted by administrating hydroxytamoxifen as described previously (15,63). Live-cell imaging showed that depletion of *Ring1a* and *Ring1b* or *Mel18* and *Bmil1* does not disperse Cbx2 condensates (**Fig. 6a**). Instead, we found that Cbx2 condensates in these double knockout mES cells are typically more and larger compared to wild-type mES cells (**Fig. 6a**). Some of Cbx2 condensates in the double knockout mES cells were irregular, instead droplet-like shape. Quantitative analysis demonstrated that both the size and the number of Cbx2 condensates in the double knockout mES cells are significantly larger than that in wild-type mES cells (**Fig. 6b-c**). These data indicate that depletion of Cbx2-PRC1 subunits increase the size and the number of Cbx2 condensates *in vivo.*

**Figure 6.**
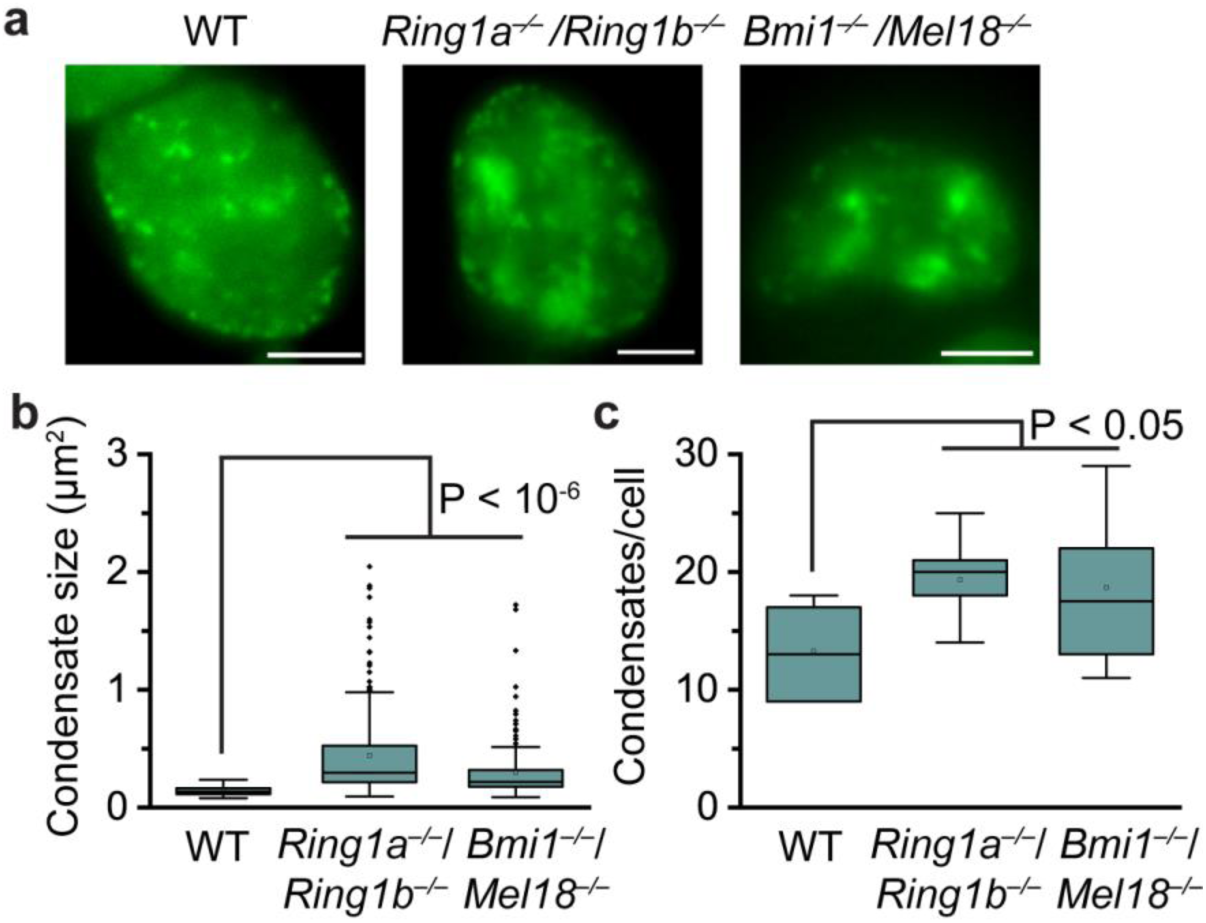
Depletion of Cbx2-PRC1 subunits facilitates the formation of Cbx2 condensate. a. Representative epi-fluorescence images for HT-Cbx2 in wild-type mES cells replicated from **Fig. 1b**, for HT-Cbx2 in *Ring1a*^−/−^*/Ring1b*^−/−^ mES cells, and for HT-Cbx2 in *Bmi1*^−/−^*/Mel18*^−/−^ mES cells. HT-Cbx2 fusions were labelled by HaloTag TMR ligand, and live-cell epifluorescence imaging was performed. Scale bar, 5.0 μm.
b. Box plot of the condensate sizes for HT-Cbx2 in wild-type mES cells replicated from **Fig. 4f**, for HT-Cbx2 in *Ring1a*^−/−^*/Ring1b*^−/−^ mES cells, and for HT-Cbx2 in *Bmi1*^−/−^*/Mel18*^−/−^ mES cells. Data were obtained from at least 10 cells. P-value was calculated based on student’s t-test.
c. Box plot of the number of condensates for HT-Cbx2 in wild-type mES cells replicated from **Fig. 4g**, for HT-Cbx2 in *Ring1a*^−/−^*/Ring1b*^−/−^ mES cells, and for HT-Cbx2 in *Bmi1*^−/−^*/Mel18*^−/−^ mES cells. Data were obtained from at least 10 cells. P-value was calculated based on student’s t-test.

## Discussion

Numerous studies have demonstrated that PcG proteins form microscopically visible condensates in primary and transformed cells, both in flies and mammals (22-26). These condensates have been shown to be the physical sites for repressing PcG-targeted genes (22-26). Consistent with these previous observations, we demonstrate that PcG protein Cbx2 forms condensates in mES cells that colocalize with H3K27me3-dense chromatin regions and Cbx2-PRC1 subunits. We further show, for the first time, that Cbx2 can undergo LLPS *in vitro* in the absence of other proteins. It is striking that Cbx2 mutants that lack the capacity of LLPS *in vitro* have a similar deficient ability to form condensates *in vivo.* The strong correlation between *in vitro* and *in vivo* data indicates that Cbx2 condensates in living cells form through LLPS. This is further supported by our following observations: Cbx2 condensates in living cells exhibit rapid exchange dynamics, a hallmark of liquid-like condensates, and are sensitive to 1,6-hexanediol, a compound known to disrupt liquid-like condensates. Our data show that Cbx2 condensates concentrate DNA and nucleosomes. Previous studies have shown that Cbx2 can compact chromatin (20). Similar to HP1α condensates compacting heterochromatin (38,39), we propose that Cbx2 compacts chromatin by forming phase-separated condensates.

The phase behavior of proteins containing IDRs can be described by the theory of associative polymers (47,64). Associative polymers phase separate through interactions between associative motifs called stickers, which are separated from one another by spacers (47,64). Spacers can impart the material properties of polymers and modulate the phase-separation ability of polymers (47,64). The stickers can be residues that involve cation-pi, electrostatic, hydrophobic, or dipolar interactions (27,28,47). In the case of Cbx2, the stickers appear to be appositively charged clusters. Perturbation of these charged clusters reduces the phase separation of Cbx2 both *in vitro* and *in vivo*, which is consistent with the notion that the phase separation of IDR-containing proteins can be promoted by interactions between blocks of appositively charged residues (39,51,54-58). The SRR also appears to be a sticker since substitution of the Ser residues of SRR with Ala prevents the phase separation of Cbx2 both *in vitro* and *in vivo.* Because the content of aromatic residues is low and there is no apparent pattern for aromatic residues across the Cbx2 sequence, we hypothesize that these aromatic residues are unlikely to be stickers. However, further experiments are required to test this hypothesis.

It has been long thought that the PRC2-catalyzed product H3K27me3 acts as a binding site for Cbx-PRC1 through its interactions with the Cbx proteins *in vivo* (61,62). This may not be the case for how Cbx2 is recruited to chromatin. We show that Cbx2 condensates can concentrate DNA and core unmodified nucleosomes and demonstrate that H3K27me3 has minimal effects on the formation of Cbx2 condensates, which is consistent with the previous observations that Cbx2 can bind DNA and compact unmodified oligonucleosomes (20,46). These results also reconcile with our live-cell single-molecule tracking results in which depletion of *Eed* or *Ezh2* has negligible effects on the chromatin bound level of Cbx2, but greatly reduces the bound level of Cbx7 and Cbx8 (15). Given that Cbx2 condensates colocalize with dense H3K27me3-marked chromatin regions, H3K27me3 could play other functions, such as organizing PcG-targeted chromatin within condensates.

Depletion of *Ring1a* and *Ring1b* or *Bmi1* and *Mel18* does not impair the formation of Cbx2 condensates, instead increases the size and the number of Cbx2 condensates. Our data also indicate that Cbx2 can form condensates in the absence of Cbx2-PRC1 subunits. Together, these results suggest that the subunits of Cbx2-PRC1 regulate the assembly and structure of Cbx2 condensates. These data can be explained by a client-scaffold model of LLPS (28,65). We propose that Cbx2 is the scaffold and the other subunits of Cbx2-PRC1 are clients. Previous studies have mapped the physical interactions between the subunits within the Cbx-PRC1 complexes. It has been suggested that one of Ring1a/Ring1b, Mel18/Bmi1, and Phc1/Phc2/Phc3 combines to form a stable heterotrimeric protein complex (66,67). The trimeric complex can be recruited into Cbx2 condensates through the C-terminal domain of Ring1b interactions with the Cbox motif of Cbx2 (68-71). It has been shown that the Phc proteins are involved in the assembly and structure of PcG condensates (23,25,26). The SAM domain can form long filaments *via* head-to-tail intermolecular interactions (72,73). We suggest two important functional roles for the trimeric client in the biogenesis and functions of Cbx2-PRC1 bodies: (1) regulating the condensate architecture, and (2) promoting proper condensate fusion by the polymerization of the Phc proteins. Thus, we propose a testable model for the assembly and structure of PcG condensates for further exploring.

In summary, our results demonstrate for the first time that PcG condensates form through LLPS and these condensates can concentrate nucleosomes and DNA. Our data show that the electrostatic interactions play a key role in promoting the phase separation of Cbx2. We further show that H3K27me3 has minimal effects on the formation of Cbx2 condensates while the Cbx2-PRC1 subunits can regulate the structure and assembly of Cbx2 condensates. Together, these results provide a starting point for conceptualizing the roles of PcG proteins in the assembly, structure, and functions of facultative heterochromatin.

## Methods

### Cell culture

PGK12.1 mES cells (74) were provided by Dr. Neil Brockdorff (University of Oxford, UK). *Cbx2*^−/−^ mES cells (75), *Eed*^−/−^ mES cells (76), *Ring1a*^−/−^*/Ring1b^fl/fl^*; *Rosa26*::CreERT2 mES cells (76), and *Bmi1*^−/−^*/Mel18*^−/−^ mES cells (76) were provided by Dr. Haruhiko Koseki (RIKEN Center for Integrative Medical Sciences, Japan). To deplete *Ring1b*, *Ring1a*^−/−^*/Ring1b^fl/fl^*; *Rosa26::CreERT2* mES cells were treated with 4-Hydroxytamoxifen (H7904, Sigma-Aldrich) for 2.0 days under a concentration of 1.0 μM as described previously (15,63). Thereafter, we referred *Ring1a*^−/−^*/Ring1b^fl/fl^*; *Rosa26::CreERT2* mES cells to *Ring1a*^−/−^*/Ring1b*^−/−^ mES cells. Dr. Tom Kerppola (University of Michigan, Ann Arbor) kindly provided HEK293T. mES cells were grown in the mES cell medium (DMEM medium (D5796, Sigma-Aldrich) supplemented with 15% FBS (97068-085, VWR), 0.1 mM non-essential amino acids (11140050, Life Technologies), 100 units per mL penicillin-streptomycin (15140-122, Life Technologies), 55 μM β-mercaptoethanol (21985-023, Life Technologies), 2 mM glutamine (G7513, Life Technologies), 10^3^ units per mL leukemia inhibitor factor) at 37 °C in 5% CO_2_. HEK293T cells were maintained in the HEK293T cell medium (DMEM (D5796; Sigma) supplemented with 10% FBS (97068-085, VWR), 100 units per mL penicillin-streptomycin (15140-122, Life Technologies), 2mM glutamine (G7513, Life Technologies), 55 μM β-mercaptoethanol (21985-023, Life Technologies)) at 37 °C in 5% CO_2_.

### Plasmids

The pTRIPZ (M1)-HT-Cbx2 plasmid harbors a puromycin-resistant gene (15). To generate Cbx2 variants fused with HT, we replaced the Cbx2 sequence in the plasmid pTRIPZ (M1)-HT-Cbx2 with the Cbx2 variant sequence. We generated the following Cbx2 variants. (1) Cbx2^ATH^, substitution of PRG with AAA (amino acid 77–79). (2) Cbx2^ATHL1^, substitution of PRG with AAA (amino acid 134–136). (3) Cbx2^ATHL2^, substitution of RKKRGRK with AAAAGAA (amino acid 161–167). (4) Cbx2^SRR^, substitution of SKSKSSSSSSSSTSSSSSS with SKSKASASASASTASASAA (amino acid 102–120). (5), Cbx2^F12A^, substitution of F-12 with A. (6) CD^Cbx2^, amino acid 1–65 of Cbx2. (7) Cbx2^ΔCD^, deletion of the CD domain (amino acid 1–65). (8) Cbx2^ΔCD-ATL^, deletion of both the CD domain and the ATL motif (amino acid 1–88).

To generate recombinant Cbx2 in *E.coli*, we amplified the Cbx2 sequence by PCR and inserted it to the downstream GST sequence within the pGEX-6P-1-GST vector (GE Healthcare, Pittsburg, PA) to generate pGEX-6P-1-GST-Cbx2. To facilitate double-affinity purification, we added a FLAG tag downstream Cbx2 sequence to generate pGEX-6P-1-GST-Cbx2-FLAG. We amplified the YFP sequence to insert the upstream Cbx2 sequence to generate pGEX-6P-1-GST-YFP-Cbx2-FLAG. To generate plasmids for expressing Cbx2 variants in *E.coli*, we amplified the sequence encoding the Cbx2 variants by PCR and used them to replace the Cbx2 sequence in the plasmid pGEX-6P-1-GST-Cbx2-FLAG. We generated the following Cbx2 variants. (1) Cbx2^ATH^, substitution of PRG with AAA (amino acid 77–79). (2) Cbx2^ATHL1^, substitution of PRG with AAA (amino acid 134–136). (3) Cbx2^ATHL2^, substitution of RKKRGRK with AAAAGAA (amino acid 161–167). (4) Cbx2^SRR^, substitution of SKSKSSSSSSSSTSSSSSS with SKSKASASASASTASASAA (amino acid 102–120).

### Establishing cell lines

24 hours before transfection, HEK293T cells were seeded into 100-mm dish to reach 85-90% of confluency at the time of transfection. Cells were co-transfected with 21 μg pTRIPZ (M) containing the fusion gene, 21 μg psPAX2, and 10.5 μg pMD2.G by using calcium phosphate precipitation. 12 hours after transfection, the medium was replaced with fresh one. 50 hours after the medium change, the medium was harvested and centrifuged at 1000 × g to remove cell debris. Cells were mixed with the harvested medium in the presence of 8.0 μg/mL polybrene (H9268, Sigma-Aldrich). 48-72 hours after transduction, 1.0–2.0μg/mL of puromycin (P8833, Sigma-Aldrich) was added to cells. Cells were selected in the presence of puromycin for at least one week.

### Generating recombinant protein of Cbx2 and its variants

Recombinant Cbx2 and its variants were generated and purified according to the previous reports (20). The pGEX-6P-1-GST-FLAG vector containing the Cbx2 fusion gene was transformed into Rosetta™ 2 (pLysS) host strains (71403, Novagen). A single colony was used to inoculate 5.0 mL of LB medium. Following overnight culture at 37 °C, 1.0 mL of the overnight culture was transferred into 1.0 L of LB. After 6-hour culture at 37 °C, the protein expression was induced overnight at 18 °C in the presence of 1.0 mM IPTG (IB02105, IBI Scientific). After centrifugation, cell pellets were resuspended in 25 mL lysis buffer (50 mM HEPES pH 7.5, 1.6 M KCl, 0.5 mM MgCl_2_, 0.5 mM EDTA, 0.06% NP-40, 1.0 mM DTT, 1.0 mg/mL lysozyme, 20 μg/mL RNase A, protease inhibitor (S8830, Sigma-Aldrich), and 0.2 mM PMSF). After three cycles of freeze-thaw by using liquid nitrogen, cells were disrupted by sonicator (VCX130, Vibra-CellTM) for 3.0 min at 45% amplitude, 15 second on, and 45 second off cycles. Cell debris was removed by centrifugation at 10,000 × g for 20 min at 4 °C. To precipitate nucleic acids, 10% polyethylenimine (PEI) in 20 mM HEPES pH 7.5 was added to the lysate to achieve a final concentration of 0.15%. The mixture was incubated for 30 min at 4 °C. After centrifugation at 20,000 × g for 20 min, supernatant was incubated with 0.5 mL of pre-washed GSH-Sepharose 4B beads (17-0756-01, GE Healthcare) for 1.0 hour at 4 °C. After washing 4 times with washing buffer (20 mM HEPES pH 7.5, 500 mM KCl, 0.2 mM EDTA, 1.0 mM DTT, and 0.2 mM PMSF), recombinant protein was eluted with 1.0 mL of 40 mM reduced L-glutathione (G4251, Sigma-Aldrich). Alternatively, recombinant protein was eluted by incubating with 80 units of PreScission protease (27-0843-01, GE Healthcare) that cleaves the GST tag at 4 °C overnight. Eluted protein was incubated with 100 μL of anti-FLAG-M2 affinity resin (A2220, Sigma) for 2.0 hours at 4 °C. After washing 4 times with washing buffer supplemented with 1.0 M KCl, recombinant protein was eluted with 0.4 mg/mL Flag peptide (F3290, Sigma-Aldrich) dissolved in washing buffer supplemented with 1.0 M KCl. Recombinant protein was resolved by SDS-PAGE to determine its purity and identity.

### *In vitro* condensate formation

After purification, KCl was added to the purified protein to reach a final concentration of 2.0 M, and then the mixture was concentrated to 30-50 μL by using Amicon centrifugal tube (UFC500324, Millipore). Protein concentration was quantified by the Bradford assay (1856209, Thermo Scientific). 30 μL of protein samples were dialyzed with Spectra/Pro 1 Dialysis Tubing (132645, Spectrum Labs) in 1.0 L of dialysis buffer (10 mM phosphate buffer pH 7.4 containing 2.7 mM KCl and 137 M NaCl) or dialysis buffer supplemented with 1.0 mM DTT at 4 °C. After changing buffer once, dialysis was performed overnight. 10 μL of the dialyzed sample was added to cover-glass dish made as described previously (77). After all condensates had settled down to the surface of coverslip, DIC or fluorescence images of condensates were acquired by using an Axio Observer D1 Microscope (Zeiss, Germany) equipped with a 100× /1.40 NA oil immersion objective with additional 2.5× magnification and an evolve EMCCD camera 512 × 512 (Photometrics, Tucson, AZ). For the excitation and emission of YFP, a Brightline^®^ single-band laser filter set (Semrock; excitation filter: FF02-482/18-25, emission filter: FF01-525/25-25, and dichroic mirror: Di02-R488-25) was used. The number of condensates per frame was counted by using ImageJ.

To determine the critical/saturation concentration of phase separation of Cbx2, a serial of concentrations of Cbx2 (1.2, 2.4, 4.8, and 12 μM) was dialyzed under the same conditions and the number of condensates per frame was counted as described above. To investigate driving forces that contribute the formation of Cbx2 condensates, to 10 μL of the dialyzed sample, NaCl, Triton X-100, and 1,6-Hexandiol (Sigma-Aldrich, 240117) was added, respectively. The mixture was incubated at 4 °C for 30 min. Condensates were imaged and analyzed as described above.

To prepare Cbx2 condensates that concentrate DNA or nucleosome, 4.8 μM of Cbx2 was mixed with Alexa 488-labelled DNA (0.5 μM), Cy5-labelled mononucleosome (40 nM), or Alexa 488 (1.0 μM), respectively. The mixture was dialyzed as described above. DIC and fluorescence images were taken by using an Axio Observer D1 Microscope as described above. For the excitation and emission of Alexa 488, a Brightline^®^ single-band laser filter set (Semrock; excitation filter: FF02-482/18-25, emission filter: FF01-525/25-25, and dichroic mirror: Di02-R488-25) was used. For the excitation and emission of Cy5, a Brightline^®^ long-pass laser filter set (Semrock; excitation filter: FF01-640/14-25, emission filter: BLP01-635R-25, and dichroic mirror: Di02-R635-25) was used. Images were processed and presented by using Photoshop.

### Live-cell imaging of condensates

Transgenic mES cells harboring HT-Cbx2 and its variants were seeded to gelatin-coated cover-glass bottom dish. After 24-hour culture at 37 °C in 5% CO_2_, cells were incubated with 5.0 nM of HaloTag^®^ TMR ligand (G8251, Promega) for 15 min at 37 °C in 5% CO_2_. After 15-min incubation, cells were washed with cell culture medium and incubated in the cell culture medium for 1.0 hour at 37 °C in 5% CO_2_. The medium was replaced with the live-cell imaging medium (FluoroBrite DMEM (A1896701), Life Technologies) supplemented with 10% FBS. Cells were maintained at 37 °C using a heater controller (TC-324, Warner Instrument) during imaging. Images of cells were acquired by using an Axio Observer D1 Microscope as described above. For the excitation and emission of TMR, a Brightline^®^ single-band laser filter set (Semrock; excitation filter: FF01-561/14, emission filter: FF01-609/54, and dichroic mirror: Di02-R561-25) was used. Visible condensates were counted by using ImageJ. Images were presented by using Photoshop.

### Live-cell imaging of YFP-Cbx2 treated with 1, 6-Hexandiol

We seeded mES cells stably expressing YFP-Cbx2 to gelatin-coated cover-glass bottom dish 24 hours before the imaging. Cell culture medium was replaced with the live-cell imaging medium and maintained at 37 °C using a heater controller. 1, 6-Hexandiol was added to the medium to reach a final concentration of 10%. Image stack was taken at every 2-min interval for 20 min by using an Axio Observer D1 Microscope as described above. For the excitation and emission of YFP, an YFP-2427B filter set (Semrock; excitation filter: FF01-500/24, emission filter: FF01-542/25, and dichroic mirror: FF520-Di02) was used.

### Immunofluorescence

Cells stably expressing *YFP*-*Cbx2* were seeded to coverslip and cultured for 24 hours. Cells were fixed with 1.0% paraformaldehyde for 10 min at room temperature. After treatment with 0.2% Triton X-100 for 10 min, cells were washed with basic buffer (10 mM PBS pH 7.2, 0.05% Tween 20, and 0.1% Triton X-100, and 0.05% Tween 20) and incubated with blocking buffer (basic buffer supplemented with 3% goat serum and 3% bovine serum albumin) overnight. Primary antibodies anti-Phc1 (39723, Active Motif, 1:200 dilution), anti Ring1B (D-319, MBL, 1:200 dilution), anti-H3K27me3 (9733, Cell Signaling Technology; 1:200 dilutions), and anti-GFP (A11122, Life Technology, 1:500 dilution) were added to cells and incubated for 2.0 hours at room temperature. After washing with the basic blocking buffer, cells were incubated with secondary antibodies Alexa 488-labeled anti–rabbit (A-11008, Life Technologies, 1:1000 dilutions) and/or Alexa 647-labelled anti-mouse (A32728, Invitrogen, 1:1000 dilution) for 2.0 hours at room temperature. Cells were washed and mounted with ProLong Antifade reagents (P7481, Life Technologies) and imaged by using Zeiss LSM 700 observer Z1 equipped with a 100× oil objective (numerical aperture, 1.4) and an electron-multiplying charge-coupled density (EMCCD) camera. For Alexa 488, 514-nm excitation and 527-nm emission filters were used. For Alexa 647, 639-nm excitation and 665-nm emission filters were used.

### FRAP

Cells stably expressing YFP-Cbx2 were seeded to 35-mm gelatin-coated cover-glass bottom dish. Cells were maintained as described for live-cell imaging. FRAP imaging was performed by using a Zeiss LSM 700 observer with the parameters: pinhole, full open; and scan speed, 177.32 (μs/pixel). Before photobleaching, two images were taken. Immediately after photobleaching, 20 images were taken with 15-s intervals. The images were analyzed using ImageJ. We used TurboReg to correct images for movement in the XY plane. After correcting fluctuations in background and total signal, the fluorescence intensities were normalized to the signal before photobleaching to obtain the fluorescence recovery as described previously (40,63).

## Acknowledgments

We thank Dr. Haruhiko Koseki for providing *Cbx2*^−/−^ mES cells, *Eed*^−/−^ mES cells, *Ring1a*^−/−^*/Ring1b^fl/fl^*; Rosa26::CreERT2 mES cells, and *Bmi1*^−/−^*/Mel18*^−/−^ mES cells and Dr. Neil Brockdorff for providing PGK12.1 mES cells. This work was supported, in whole or in part, by the National Cancer Institute under Award Number R03CA191443 (to X.R.), the National Science Foundation under Award Number CHE-1500285 (to H.W.), the National Institute of General Medical Sciences under Award Number R01GM098401 (to T.Y.), and the Office of Research Services (ORS) at University of Colorado Denver.

## Author contributions

X.R. conceived and designed the study, supervised and performed the experiments, analyzed data, prepared the figures, and wrote the paper. R.T. established transgenic mESC lines, performed *in vitro* condensate assay, live-cell imaging, immunofluorescence, and analyzed data. S.K. constructed plasmids and analyzed data. K.B. constructed plasmids. Y.T. provided reconstituted nucleosomes. H.W. performed data analysis and supervised students. T.N.H., H.N.D., and C.Z. constructed plasmids and established transgenic mESC lines. B.M. analyzed data. All authors provide constructive comments for the manuscript.

